# Generation of four iPSC lines from peripheral blood mononuclear cells (PBMCs) of an Attention Deficit Hyperactivity Disorder (ADHD) individual and a healthy sibling in an Australia-Caucasian family

**DOI:** 10.1101/420505

**Authors:** Janette Tong, Kyung Min Lee, Xiaodong Liu, Christian M Nefzger, Prasidhee Vijayakumar, Ziarih Hawi, Ken C Pang, Clare L Parish, Jose M Polo, Mark A Bellgrove

**Affiliations:** Monash Institute of Cognitive and Clinical Neurosciences and School of Psychological Sciences, Monash University, Wellington Road, Clayton, Victoria, Australia; Department of Anatomy and Developmental Biology, Monash University, Wellington Road, Clayton, Victoria, Australia; Development and Stem Cells Program, Monash Biomedicine Discovery Institute, Wellington Road, Clayton, Victoria, Australia; Australian Regenerative Medicine Institute, Monash University, Wellington Road, Clayton, Victoria, Australia; Department of Genetic Engineering, School of Bioengineering, SRM Institute of Science and Technology, Kattankulathur 603203, India; Murdoch Children’s Research Institute, Parkville, VIC, Australia; Royal Children’s Hospital, Melbourne, VIC, Australia; Department of Pediatrics, University of Melbourne, Parkville, VIC, Australia; Department of Psychiatry, University of Melbourne, Parkville, VIC, Australia; Florey Institute of Neuroscience and Mental Health, The University of Melbourne, Parkville, VIC 3010, Australia

## Abstract

Peripheral blood mononuclear cells were donated by a male teenager with clinically diagnosed Attention Deficit Hyperactivity Disorder under the Diagnostic and Statistical Manual of Mental Disorders IV criteria and his unaffected male sibling. Induced pluripotent stem cells were developed using integration-free Sendai Reprogramming factors containing OCT, SOX2, KLF4, and c-MYC. All four iPSC lines displayed pluripotent cell morphology, pluripotency-associated factors at protein level, alkaline phosphatase enzymatic activity, male karyotype of 46, XY, and *in vitro* differentiation capacity into all the three germ layers and negative for Mycoplasma.

## Introduction

The pathophysiology of Attention Deficit Hyperactivity Disorder (ADHD) is poorly understood due to the lack of cellular models that faithfully represent the clinical ADHD features, and limited live tissues to expand for long-term mechanistic studies. We generated iPSCs from an ADHD patient and a unaffected sibling to understand the pathophysiology.

## Results

A blood sample was donated from a pair of dizygotic twin brothers with one brother meeting the Diagnostic and Statistical Manual of Mental Disorders IV (*DSM-IV*) criteria (DuPaul et al., 2001) for ADHD and his male sibling being unaffected. Both individuals completed the Conners’ Parent Rating Scale-Revised: Long Form (CPRS-R:L) which is designed to assess symptoms (inattention, impulsivity, hyperactivity) of ADHD (Conners et al., 1998). For the child’s gender and age, the affected sibling scored above the 98^th^ percentile on the Impulsivity subscale and above the 95^th^ percentile on the Hyperactivity, ADHD Index and *DSM-IV* Inattentive subscales of CPRS-R:L, which is consistent with the diagnostic interview conducted by an experienced psychiatrist. The unaffected sibling scored at or below average *T*-Scores and percentiles across all CPRS-R:L subscales. The affected sibling is managed by Ritalin and fluoxetine at the time of testing and sample collection. The study was approved by the Monash University Human Research Ethics Committee Australia under the approval number, CF15/2566 – 2015001048.

Four induced pluripotent stem cell (iPSC) lines (two from the affected sibling-MICCNi002-A and –B; two from the unaffected sibling - MICCNi001-A and –B) were generated from peripheral blood mononuclear cells (PBMCs) by reprograming with integration-free Sendai viral vectors expressing *KLF4, OCT3/4, c-MYC* and *SOX2*. The PBMCs were initially seeded in PBMC medium and half of the medium was replaced with fresh medium until the transduction, where the reprogrammed cells were plated on irradiated Mouse Embryonic Fibroblasts (MEFs) the subsequent day (Nefzger et al., 2014). The cells were grown on MEFs for 20 days with an addition of fresh MEF culture dishes every 7 days to allow colony formation, and large colonies were dissected and transferred to vitronectin-coated flasks. All iPSC colonies showed a round compact shape with clear peripheral outline, which is typical colony morphology (Fig. 1A). RT-PCR analyses of the viral genome and transgenes confirmed that our iPSC lines were Sendai-negative and vector-free after 10 passages (Fig. 1B).

Immunocytochemistry analyses of MICCNi001-A and –B and MICCNi002-A and –B at passage number 12 showed strong expression of pluripotency markers TRA 1-60, OCT4, NANOG and KLF4, as well as positive for alkaline phosphatase enzymatic activity (Fig. 1C and 1D). Mycoplasma testing of all lines at passage 15 was negative by luminescence (Supp. File 1). G-band analyses at passage 10 exhibited normal male karyotypes of 46, XY for MICCNi002-A (Fig. 1E) and others (Suppl. File 1). Our iPSC lines have adequate potential for *in vitro* differentiation into the three germ layers via embryoid bodies using a conventional method (Thermofisher, 2013) (Fig. 1F). Subsequently, the generated embryoid bodies were tested for expression of >80 differentiation associated markers for all three germ layers as well as key pluripotency genes using the hPSC Scorecard assay. All *in vitro* differentiated iPSC lines showed reduced expression of self-renewal genes and upregulation of markers of mesendoderm, ectoderm, mesoderm and endoderm (Fig 1G). Lastly, MICCNi001-A, MICCNi001-B, MICCNi002-A and MICCNi002-B lines were authenticated with 100% concordance with its parental PBMCs using Short Tandem Repeat profiling (Supp. File 2).

## Materials and Methods

### 1. Human sample

Human peripheral blood mononuclear cells (PBMCs) were isolated from 30 mL blood in Leucosep™ Tubes (Greiner Bio-One) filled with 15 mL Ficoll-Paque Plus^™^ (GE Healthcare). Within 1 hour of collection, the blood sample was centrifuged at 800 × *g* for 15 min at room temperature (RT), washed three times with PBS and centrifuged at 250 × *g* for 10 min at RT. 1 × 10^6^ cells were frozen in 20% Dimethyl sulfoxide and 80% Fetal Bovine Serum (Bovogen) until Sendai transduction.

### 2. PBMC reprogramming, iPSC generation and culture

After recovery from cryopreservation PBMCs (5 × 10^5^ cells per well of a 24-well plate 4 days prior to transduction) were cultured in PBMC medium (Thermofisher, 2016) until transduction with Sendai virus particles containing *OCT, SOX2, KLF4*, and *c-MYC* using CytoTune^®^-iPS 2.0 Sendai Reprogramming Kit (Life Technologies). Transduced cells were cultured on growth inactivated/irradiated mouse embryonic fibroblasts (MEFs) feeder-cells for 7 days in StemPro^®^-34 medium without cytokines. Thereafter cells were fed with fresh iPSC medium (Thermofisher, 2016) and monitored daily until large colonies were formed within a two week period. Each colony was manually picked, transitioned in culture vessels coated with Vitronectin (Thermofisher) and expanded in E8 medium (Invitrogen).

### 3. Immunocytochemistry and Alkaline Phosphatase staining of iPSC

Pluripotency of iPSCs cultured on Vitronectin-coated plates were examined by immunocytochemistry analyses with TRA 1-60, OCT3/4, NANOG or KLF4 antibodies. Cells were seeded at 1 × 10^5^ cells on an eight-chamber, polystyrene vessel tissue culture treated glass slide (Falcon, USA) for 6 days. Cells were fixed in 4% formaldehyde solution (Merck) for 15 min at RT, washed with PBS and incubated with 0.1% Triton X 100 for 30 min. Cells were subsequently washed with PBS and incubated with antibodies for TRA 1-60, OCT3/4, NANOG or KLF4 (Table 3) for 24 hours at 4°C. Cells were then washed three times with PBS and incubated with Secondary Alexa Fluor antibodies (Table 3) for 2 hours. Lastly, cells were washed 3 × PBS and fixed with Fluroshield^™^ with DAPI (Sigma-Aldrich). Images were acquired using IX71 inverted microscope (Olympus) and processed in ImageJ software. Alkaline phosphatase staining was performed using Vector^®^ Black Alkaline Phosphatase Substrate Kit II as per manufactures instructions (Vector Laboratories, USA).

**Table 1:** Summary of lines

| iPSC line names | Abbreviation in figures | Gender | Age | Ethnicity | Genotype of locus | Disease |
| --- | --- | --- | --- | --- | --- | --- |
| MICCNi001-A | MICCNi001-A | Male | 16 | Caucasian | N/A | Healthy |
| MICCNi001-B | MICCNi001-B | Male | 16 | Caucasian | N/A | Healthy |
| MICCNi002-A | MICCNi002-A | Male | 16 | Caucasian | N/A | Attention Deficit Hyperactivity Disorder |
| MICCNi002-B | MICCNi002-B | Male | 16 | Caucasian | N/A | Attention Deficit Hyperactivity Disorder |

**Table 2:**
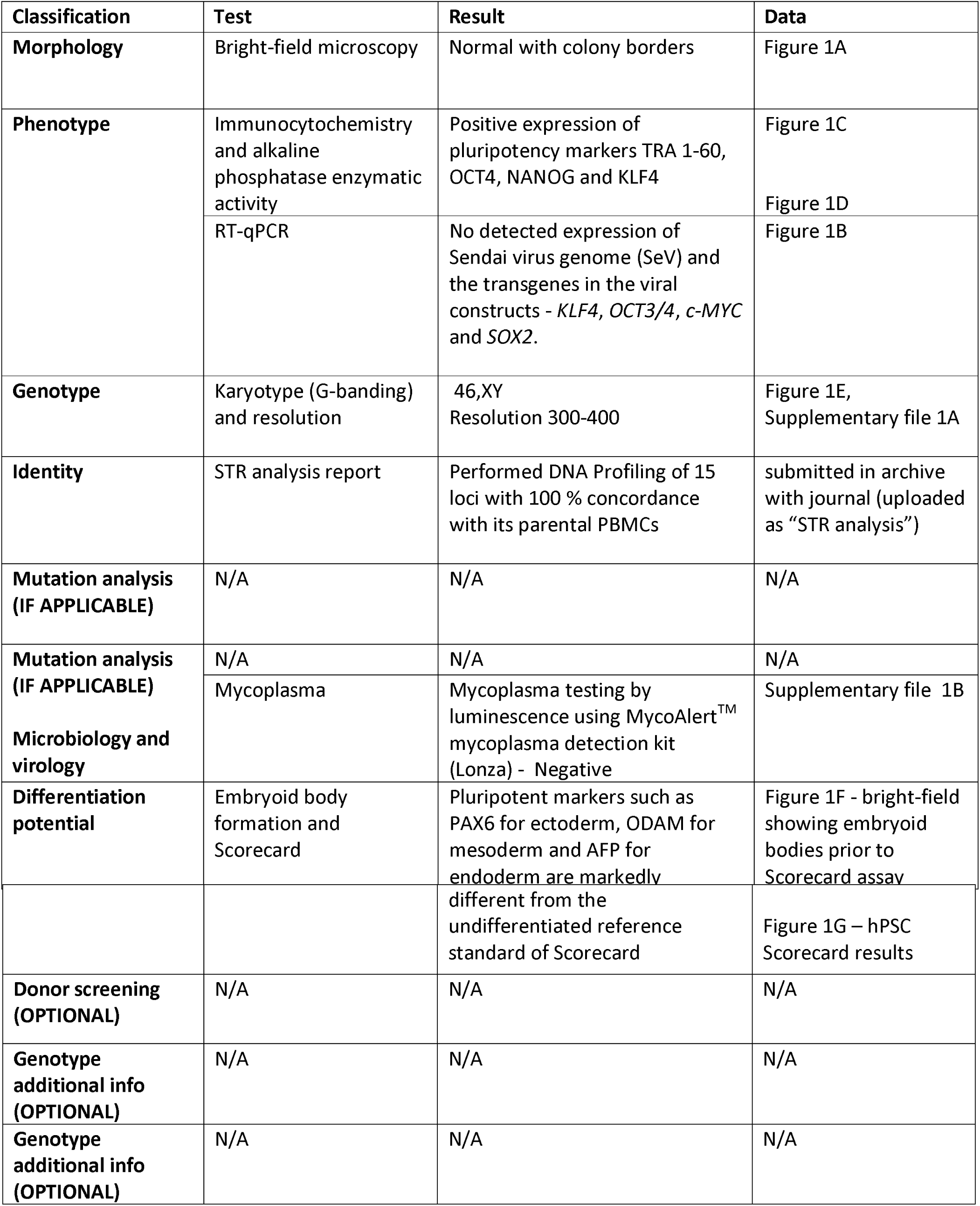
Characterization and validation

**Table 3:**
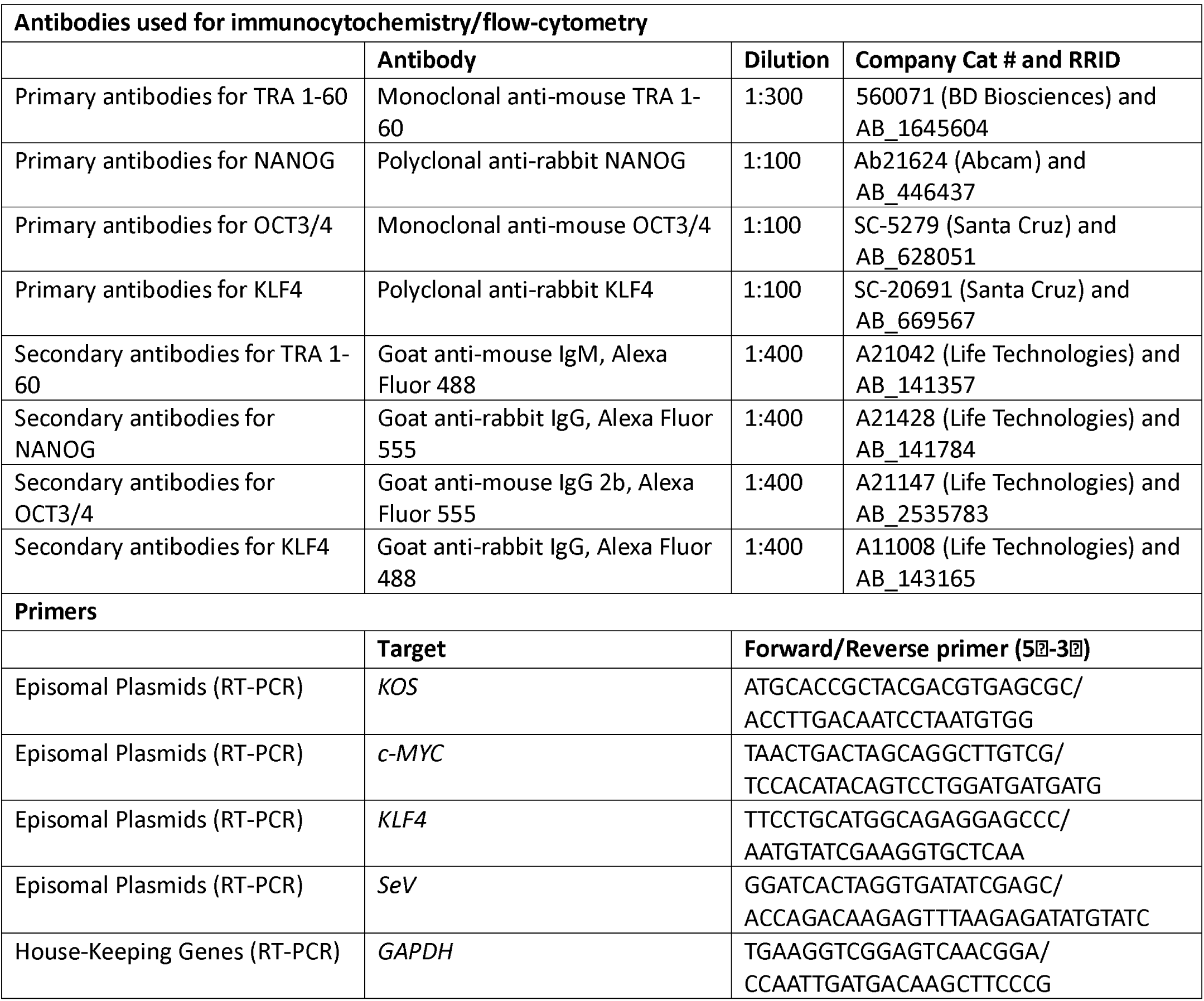
Reagents details

### 4. Karyotyping

G-banding of iPSCs was carried out at the Monash Pathology Services (Monash Medical Centre).

### 5. Embryoid bodies (EB) formation and ScoreCard

The differentiation capacity of iPSC lines was analyzed by the formation of EB as per manufacturer’s instructions of the ScoreCard assay (Thermofisher). Briefly, iPSCs were dissociated and culture non-adherently in EB differentiation medium for 7 days. Total RNA of cells were extracted using the RNeasy micro kit (Qiagen) and reverse transcribed into cDNA using the SuperScript III cDNA synthesis kit (Invitrogen). Quantitative analyses of cDNA samples were performed using the TaqMan hPSC Scorecard Assay (Thermofisher).

### 6. Short Tandem Repeat analysis

DNA extracted from PBMCs and iPSCs were sent to the Medical Genomics Facility (MHTP, Melbourne) for STR analysis, where 16 loci were investigated by PowerPlex HS16 System kit (Promega).

## Acknowledgements

We acknowledge the financial support of the National Health and Medical Research Council [Early Career Fellowship ID 1112452 (JT)], the Society for Mental Health Research [Early Career Research Project Grant Award (JT)], the Rebecca L Cooper Medical Research Foundation [Medical Research Grant ID 10409 (JT)] and Monash University [Strategic Grant Scheme ID SGS16-0410 (JT)].

---

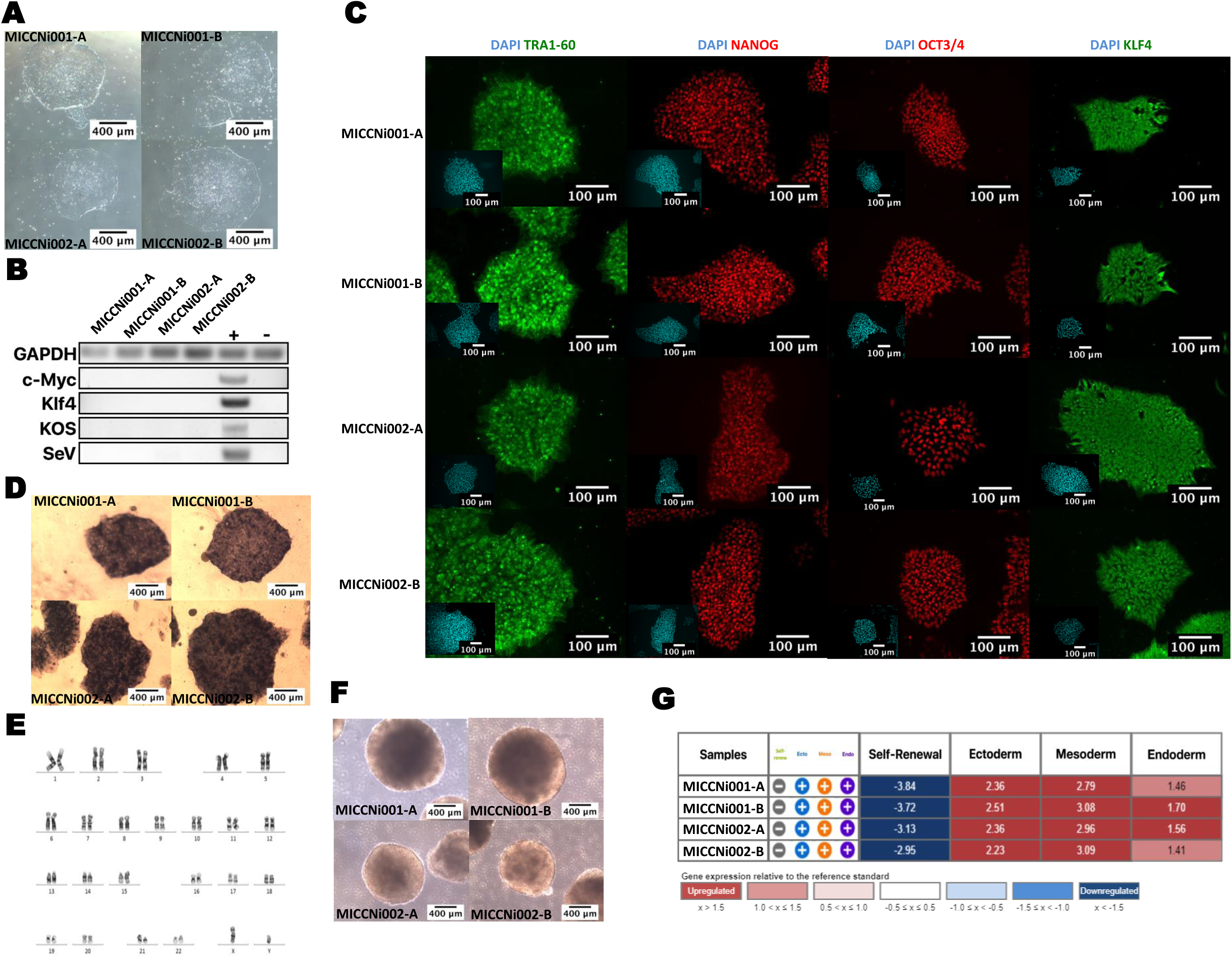

